# Mature microRNA-binding protein QKI promotes microRNA-mediated gene silencing

**DOI:** 10.1101/2023.03.15.532749

**Authors:** Kyung-Won Min, Myung Hyun Jo, Minsuk Song, Seungbeom Ko, Ji Won Lee, Min Ji Shim, Kyungmin Kim, Hyun Bong Park, Shinwon Ha, Hyejin Mun, Ahsan Polash, Markus Hafner, Jung-Hyun Cho, Dong-San Kim, Sungchul Hohng, Sung-Ung Kang, Je-Hyun Yoon

## Abstract

Although Argonaute (AGO) proteins have been the focus of microRNA (miRNA) studies, we observed AGO-free mature miRNAs directly interacting with RNA-binding proteins, implying the sophisticated nature of fine-tuning gene regulation by miRNAs. To investigate microRNA-binding proteins (miRBPs) globally, we analyzed PAR-CLIP data sets to identify RBP quaking (QKI) as a novel miRBP for let-7b. Potential existence of AGO-free miRNAs were further verified in genetically engineered AGO-depleted human and mouse cells. We have shown that QKI serves as an auxiliary factor empowering AGO2/let-7b-mediated gene silencing. Depletion of QKI decreases interaction of AGO2 with let-7b and target mRNA, consequently controlling target mRNA decay. QKI, however, also suppresses the dissociation of let-7b from AGO2, and slows assembly of AGO2/miRNA/target mRNA complexes at the single-molecule level. We also revealed that QKI suppresses cMYC expression at post-transcriptional level, and decreases proliferation and migration of HeLa cells, demonstrating that QKI is a tumor suppressor gene by in part augmenting let-7b activity. Our data show that QKI is a new type of RBP implicated in the versatile regulation of miRNA-mediated gene silencing.

## INTRODUCTION

Post-transcriptional gene regulation is a key mechanism for eukaryotic organisms, enabling them to respond rapidly to environmental changes by synthesizing proteins with proper timing and location (Gehring *et al*, 2017; Jackson *et al*, 2010; Jonas & Izaurralde, 2015). This regulatory mechanism is mainly governed by three major factors: RNA-binding proteins (RBPs), non-coding RNAs such as microRNAs (miRNAs), and long non-coding RNAs (lncRNAs) (Gehring *et al*., 2017; Ha & Kim, 2014; Yoon *et al*, 2013a). Approximately 2,300 or higher number of miRNAs are expressed in human cells, which target complementary sequences in mRNAs and lncRNAs, to suppress mRNA translation or induce target RNA degradation (Alles *et al*, 2019; Chiang *et al*, 2010; Ha & Kim, 2014). The mechanism(s) of miRNA biogenesis and gene silencing have been well studied whereas miRNA decay occurs via terminal modification and target-directed miRNA degradation (Chatterjee & Großhans, 2009; Han *et al*, 2020; Shi *et al*, 2020; Towler *et al*, 2015).

In order for miRNAs to silence target mRNAs, mammalian cells require specialized proteins to promote target mRNA decay and suppress translation. Argonaute (AGO) proteins are direct binding partners of mature miRNAs, and central components of the RNA-induced Silencing Complex (RISC). DICER1 directly interacts with the AGOs and transfers a miRNA duplex to AGOs, with the thermodynamically favored guide strand being retained, and the passenger strand being discarded from the AGOs (Kawamata *et al*, 2009; Kim *et al*, 2016). The AGO-bound miRNAs then guide the AGOs to target mRNAs for decapping, deadenylation, and exo-ribonucleolytic decay (Fabian & Sonenberg, 2012).

Recently, miRNA’s fate after AGO-mediated gene silencing has been explored using RNA labeling and high throughput RNA-seq analysis (Baccarini *et al*, 2011; Becker *et al*, 2019; Elbarbary *et al*, 2017; Ghini *et al*, 2018; Kingston & Bartel, 2019; Kleaveland *et al*, 2018; Marzi *et al*, 2016; Sheu-Gruttadauria *et al*, 2019; Shukla *et al*, 2019). The most mature miRNAs are highly stable (Reichholf *et al*, 2019) and are produced at rates as fast as 110 ± 50 copies/cell/min in mouse embryonic fibroblasts, and miRNAs spanning a highly complementary target site are prone to be degraded for AGO2 recycling since AGO2 outlives miRNAs (Kingston & Bartel, 2019). Decay of selective mature miRNAs is either influenced by the 3’-end trimming of pre-miRNAs and mature miRNAs (Katoh *et al*, 2009; Lee *et al*, 2019; Li *et al*, 2005; Shukla *et al*., 2019) or by proteasome-dependent turnover of AGO2 when miRNA-mRNA interacts via extensive base paring (Han *et al*., 2020; Shi *et al*., 2020).

In this study, our findings suggest that several types of RBPs, including quaking (QKI), directly bind a subset of mature miRNAs. QKI was previously implicated in mRNA splicing, export, and decay (De Bruin *et al*, 2016; Larocque *et al*, 2002; Thangaraj *et al*, 2017) and circular RNA biogenesis (Conn *et al*, 2015). Specifically, our observations indicated that QKI: (i) directly binds a subset of mature miRNAs, (ii) promotes the interaction of AGO2 with let-7b, (iii) decreases the stability of mRNA targeted by let-7b, (iv) inhibits the dissociation of let-7b from AGO2, and (v) delays the kinetics of AGO2/miRNA/target mRNA complex assembly at a single-molecular level. Furthermore, our results also suggest that QKI imparts anti-tumorigenic effects partly by enhancing AGO2/let-7b/*cMYC* axis. Here, we reveal newly identified QKI as a novel miRNA binding protein and add an additional layer of complexity in miRNA-mediated gene silencing.

## RESULTS

### Canonical RBPs directly bind mature miRNAs, which are free of AGO proteins

In order to profile RBPs directly binding with mature miRNAs, we first analyzed the published CLIP data sets (2075 in total) and then searched RBPs (311 in total) cross-linked with mature miRNA let-7 family, miR-21, and miR-130b (Zhu *et al*, 2019). Since Photoactivatable Ribonucleoside-Enhanced Crosslinking and Immunoprecipitation (PAR-CLIP) data provides the most reliable measure of RBP and RNA crosslinking based on T-to-C conversion (Hafner *et al*, 2010), we plotted genomic coordinates of let-7a-1, let-7b, let-7c (**Figure 1** and **Table S1**), let-7a-2, let-7a-3, let-7d (**Figure S1** and **Table S1**), miR-21 and miR-130b (**Figure S2** and **Table S1**) corresponding to their mature miRNA sequences. We chose those miRNAs as candidates because they are well-characterized in cellular processes and human diseases (Büssing *et al*, 2008; Massey *et al*, 2018; Sheedy, 2015; Wang *et al*, 2010b; Yamamichi *et al*, 2009; Zhang *et al*, 2016). Our analysis revealed 7 RBPs binding with mature let-7a-1 (**Figure 1A**), 36 RBPs with let-7b (**Figure 1B**), 29 RBPs with let-7c (**Figure 1C**), 16 RBPs with let-7d (**Figure S1B**), 87 RBPs with miR-21 (**Figure S2A**), and 37 RBPs with miR-130b (**Figure S2B**). T-to-C conversion ratio also indicates that the binding with RBPs and mature miRNA is comparable to AGO binding (**Figure 1** and **Figure S1-S2**). Common RBPs binding with let-7, miR-21 and miR-130b include AGO, HuR, AUF1 and QKI (**Figure S1C and S2C**). These results demonstrate that there is a pool of RBPs directly binding with mature miRNAs.

**Figure 1.**
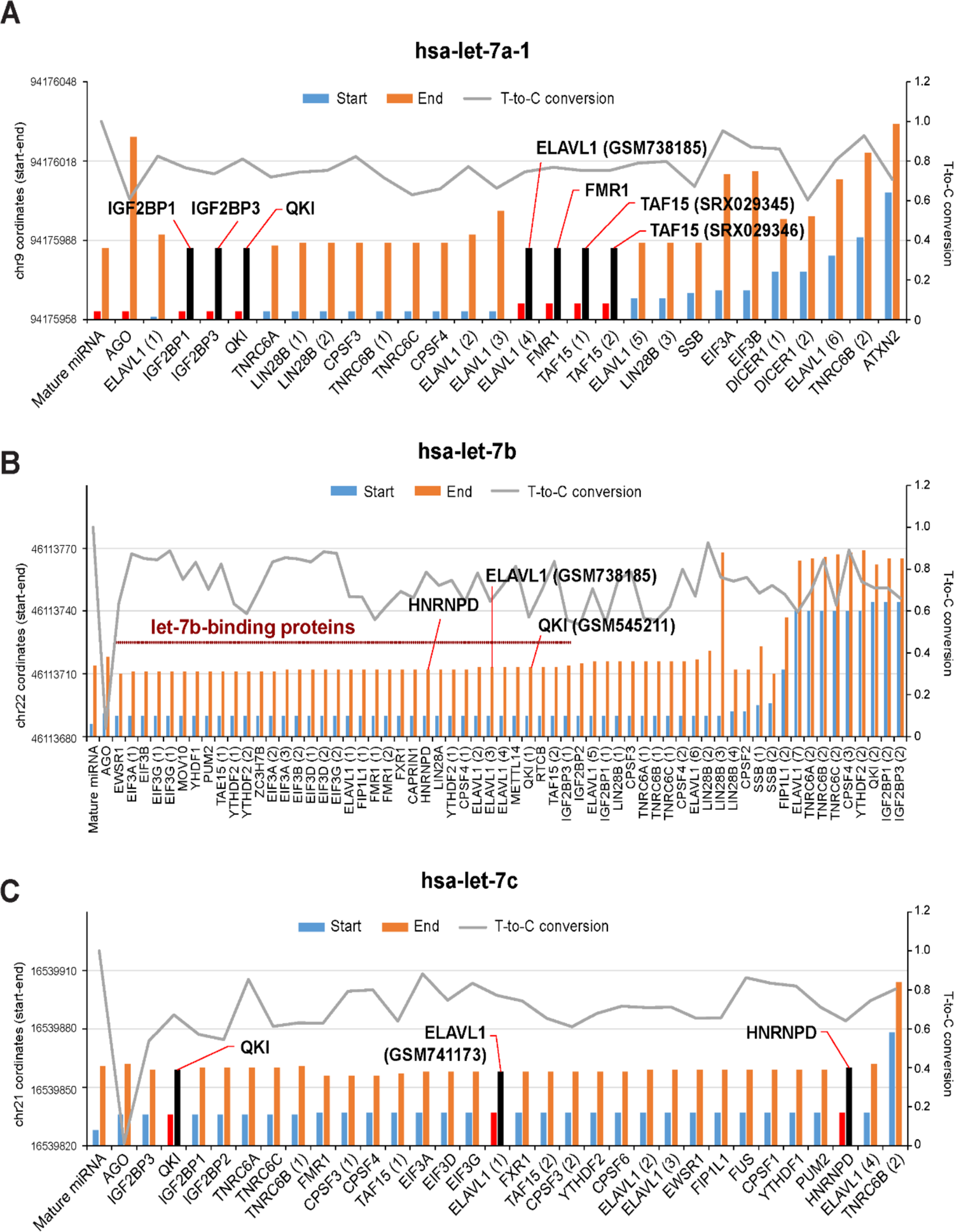
PAR-CLIP analysis identified RNA-binding proteins directly binding with mature and precursor miRNAs. **(A - C)** Genome coordinates (y-axis) of precursor and mature miRNA let-7a-1 (A), 7b (B), and 7c (C) identified from PAR-CLIP using human cell lines. T-to-C conversion ratio was presented in y-axis as well. Representative RBPs such as QKI, HuR, hnRNPD/AUF1 and IGF2BP were highlighted.

MiRNAs are processed by DROSHA and DICER, and then loaded directly to AGO proteins (Gregory *et al*, 2004; Hafner *et al*., 2010; Lee *et al*, 2003). There are many reports that mature miRNAs generally are degraded rapidly after dissociation with AGOs (Kingston & Bartel, 2019; Reichholf *et al*., 2019). Since our findings in **Figure 1** demonstrated that many RBPs directly bind with mature miRNAs, one possibility is that there is a pool of mature miRNAs resistant to degradation after dissociation from AGO proteins. To prove this idea, we set up the assay of AGO immunodepletion using antibodies recognizing AGO2 or Pan-AGO proteins. After series of immunoprecipitation using human HeLa cells and mouse embryonic fibroblasts (MEFs) (**Figure 2A and 2B**), we measured amount of let-7b, miR-21, and miR-130b from total RNAs or supernatants after the third immunoprecipitation. Our RNA-seq data revealed that ∼10% of mature let-7b, let-7i and miR-21 still exist in the lysates after immunodepletion of AGO2 and Pan-AGO in human cells (**Figure 2C** and **Table S2**). However, let-7c, let-7e, and miR-130b were depleted in the supernatant after the third immunoprecipitation suggesting that they are quickly degraded after dissociation from AGO proteins (**Figure 2C**). In contrast, mature miRNAs from MEFs are still observed after AGO2 or Pan-AGO depletion (∼10%) indicating that mouse cells contain more mature miRNAs free of AGO proteins (**Figure 2D** and **Table S2**). These results indicate that a subset of mature miRNAs exist in the form of AGO-free complex.

**Figure 2.**
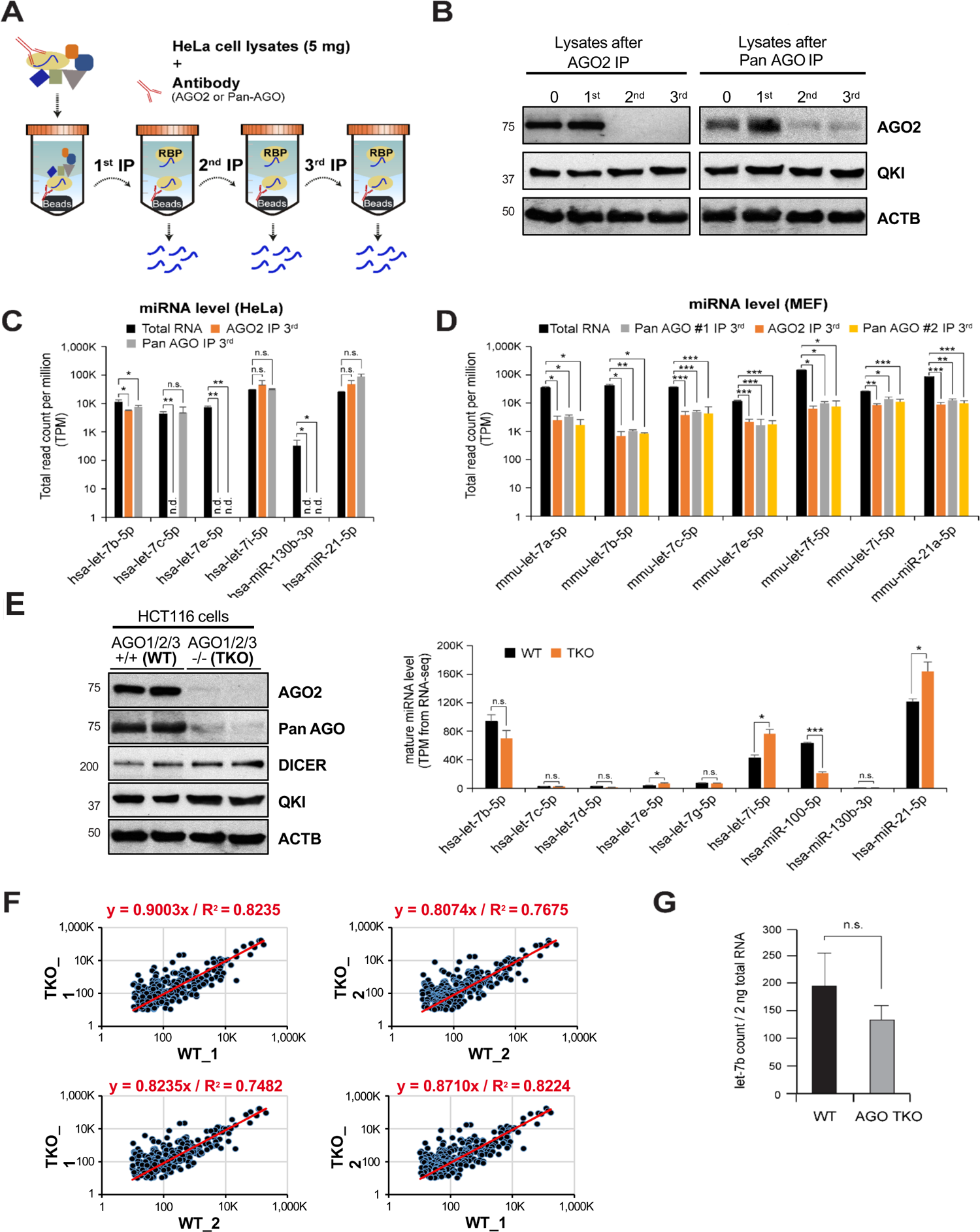
Existence of AGO-free mature miRNAs in human and mouse cells. **(A)** Schematic of experimental set up to profile mature miRNAs from AGO-depleted human and mouse cell lysates. HeLa or MEF lysates were immunoprecipitated using anti-AGO2 or anti-Pan AGO antibodies serially. The resulting pellets and lysates were subjected to RNA purification followed by high throughput RNA-sequencing or RT-qPCR. **(B)** Western blot analysis of AGO2, QKI and ACTB using cell lysates before and after serial immunoprecipitation. **(C, D)** RNA-seq analysis of let-7, miR-21 and miR-130b using cell lysates before and after immunoprecipitation with anti-AGO2 or anti-Pan AGO antibodies. N = 2-4, n.d., not detected from RNA-seq, \*\*\**p* < 0.001, \*\**p* < 0.01, \**p* < 0.05, *n.s*., not significant from Student’s *t*-test. Data are mean ± S.D. **(E)** Western blot analysis of AGO2, Pan-AGO, DICER, QKI and ACTB using cell lysates HCT116 cells targeting AGO1, 2, and 3. RNA-seq analysis of let-7, miR-21, miR-130b, and miR-100 using HCT116 wild type and AGO TKO cells. N = 2, \*\*\**p* < 0.001, \**p* < 0.05, *n.s.*, not significant from Student’s *t*-test. Data are mean ± S.D. **(F)** Scatter plots of miRNAs sequenced from HCT116 wild type and AGO TKO cells. **(G)** Counting of let-7b from 2 ng total RNA purified from HCT116 wild type and AGO TKO cells. N = 6, *n.s.*, not significant from Student’s *t*-test. Data are mean ± S.D.

Genetic deletion of either AGO1 or AGO2 as well as double deletion marginally affects level of mature let-7b both in miRNA counting and RNA-seq (**Figure S3A**). Triple knockout of AGO1, 2 and 3 in HCT116 cells (AGO TKO) also revealed that there is still significant amount of mature let-7b, miR-21, miR-100, and let-7i even when there are no AGO proteins (**Figure 2E *Right* and Table S2**). RNA-seq data also supported our findings that genetic deletion of AGO1, 2, and 3 did not affect mature miRNA level globally (**Figure 2F**) as observed in *Dicer* and *Drosha* knockout (Ruby *et al*, 2007; Yang *et al*, 2010). Although the expression of passenger strand increased in AGO TKO cells, the mature strands were predominantly expressed in AGO TKO cells (**Figure S3B and Table S2**). We also confirmed that comparable let-7b copy numbers are presented in AGO TKO cells compared with WT using AGO-FISH assay (**Figure 2G**). In MEFs, level of let-7b was resistant to deletion of AGO2 both in RNA-seq and miRNA counting (**Figure 2D and Figure S3C**), and even the stability of miRNAs did not change significantly (**Figure S3D**). These findings demonstrate that there are a portion of mature miRNAs which are not associated with AGO proteins and suggest that there should be distinct function of RBPs directly binding with mature miRNAs.

### QKI enhances interaction of AGO2 with let-7b

Since our results showed that QKI is associated with let-7 family (**Figure S1C**), we thereafter focused on let-7b, mainly due to its known functions in normal physiology and pathogenesis such as cancer, for investigating QKI’s role in miRNA function. Direct interaction of QKI and let-7b predicts competition of QKI with AGO2 for let-7b binding (**Figure 3A**). To test this hypothesis, we depleted QKI in HeLa cells and, using Ribonucleoprotein Immunoprecipitation (RIP) RT-qPCR analysis, determined how this depletion affects the interaction of AGO2 with let-7b. Our analysis showed that depletion of QKI by short hairpin RNA (shRNA) significantly decreased the abundance of let-7b in AGO2 immunoprecipitates without affecting the expression of let-7b (**Figure 3B**). These results suggest that QKI promotes interaction of AGO2 and let-7b in HeLa cells.

**Figure 3.**
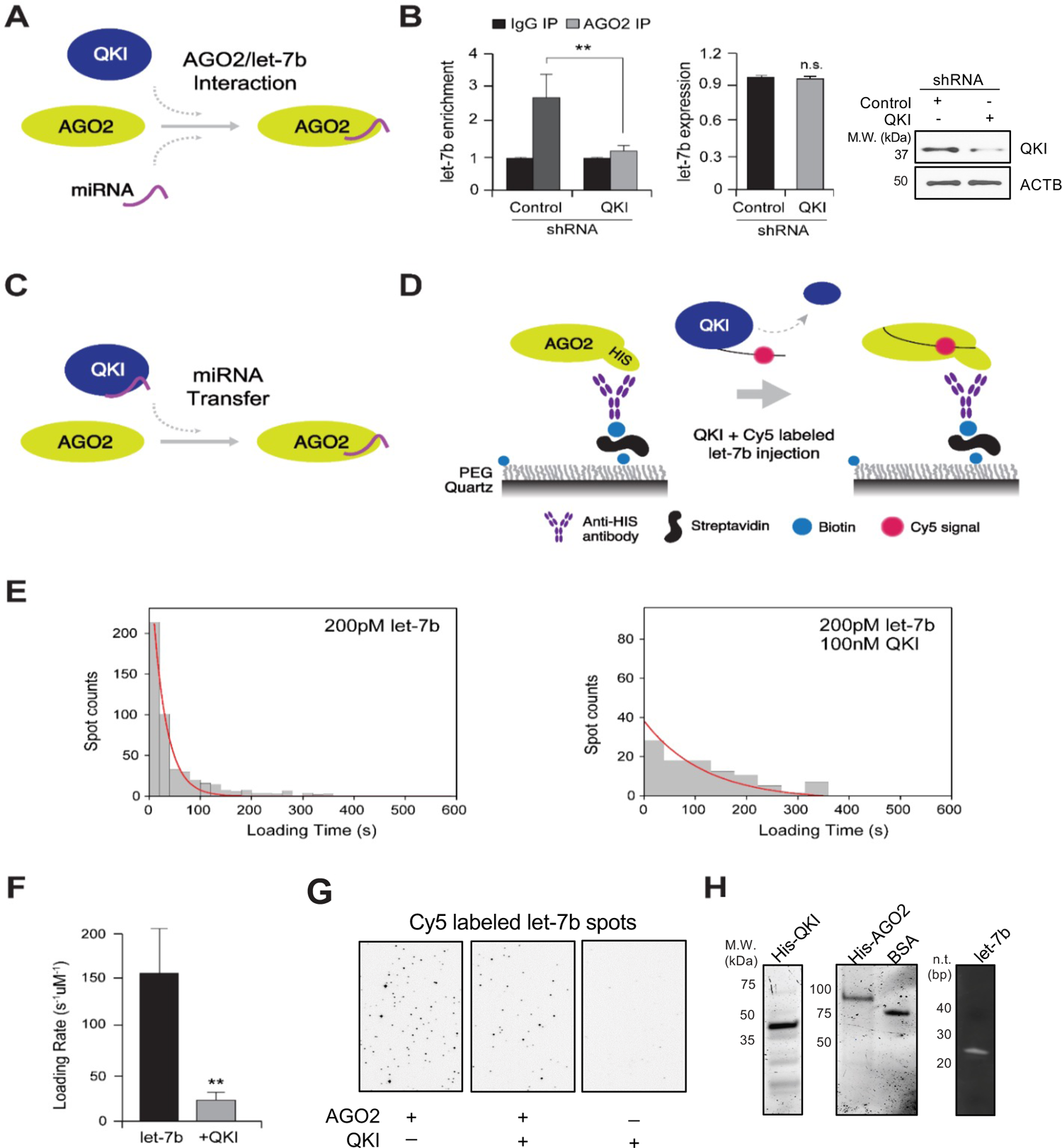
QKI is required for AGO2 interaction with let-7b and allows let-7b loading onto AGO2. (A) A model of QKI’s effect on interaction of AGO2 and let-7b in human cells. (B) AGO2 RIP assay in HeLa cells after transfection of control and QKI shRNA. (*Left*) At 48 hours after transfection of shRNA targeting GFP or QKI in HeLa cells, cell lysates were subjected to immunoprecipitation against AGO2 for detection of co-purified let-7b by RT-qPCR. (*Middle*) At 48 hours after transfection of shRNA targeting GFP or QKI in HeLa cells, total RNAs were purified for detection of let-7b by RT-qPCR. (*Right*) Efficiency of QKI shRNA was determined by western blot of total cell lysates with anti-QKI or anti-ACTB antibodies. N = 3, \*\**p* < 0.01, *n.s.*, not significant from Student’s *t*-test. Data are mean ± S.D. Data are mean ± S.D. (C) A model of QKI’s effect on let-7b loading on AGO2 in single molecule analysis. (D) Single-molecule experimental scheme for let-7b (guide RNA) loading on AGO2. Recombinant AGO2 was immobilized on PEG-passivated surface using avidin-biotin interaction and anti-His tag antibody in a detection chamber. Cy5-labeled let-7b was injected into the chamber with or without QKI during the imaging. (E) Histograms of let-7b loading times in (*Left*) absence or (*Right*) presence of recombinant OKI. The red lines are single exponential decay fit (27 s and 130 s, respectively). (F) Single exponential fit was performed for loading time distributions. Loading rates of let-7b onto immobilized AGO2 with or without QKI. About 300-500 loading events were observed per experiment. N = 3, \*\**p* < 0.01, from Student’s *t*-test. Data are mean ± S.D. (G) Cy5-labeled let-7b spots on slides with or without immobilized AGO2 and QKI 5 minutes after let-7b injection. The results of single-molecule assay. (H) Recombinant QKI and AGO2 were visualized by Coomassie Blue staining and let-7b is imaged by ethidium bromide staining. The abundance of let-7b was assessed by reverse transcriptase PCR analysis.

To address how QKI facilitates assembly of AGO2-let-7b complexes, we investigated miRNA transfer (**Figure 3C**) by performing the single-molecule AGO2 protein-miRNA binding assay as we reported previously (Yoon *et al*, 2015). To observe let-7b loading onto AGO2 in real-time, recombinant human AGO2 proteins were immobilized on a PEG-passivated and biotin-functionalized surface using an anti-His tag antibody (**Figure 3D**). Cy5-labeled let-7b (200 pM) was injected with or without recombinant QKI (100 nM) and the appearance of let-7b fluorescence signals were monitored. Even when let-7b was pre-bound to QKI, we observed loading, showing that the let-7b can be transferred from QKI to AGO2. The let-7b loading rate with QKI (**Figure 3E-H**) was slower than that with AUF1 previously reported (Yoon *et al*., 2015). These findings suggest that QKI facilitates the assembly of AGO2-let-7b in cells by directly transferring let-7b to AGO2. It also implicates that transient formation of a QKI:AGO2:let-7b ternary complex slows both on- and off-rates of miRNA. If the effect of QKI on the let-7b off-rate is significantly more than on the let-7b on-rate, the steady-state equilibrium would be pushed in favor of the let-7b-bound complex, as reflected in the RIP data. (**Figure 3B, and see discussion**).

### QKI promotes decay of let-7 target mRNAs

Next, we tested the consequence of QKI’s regulation of AGO2 binding to let-7b on target mRNA decay. We compared enrichment of a let-7b target mRNA, *POLR2D* mRNA, as predicted in TargetScan and in our previous study (Yoon *et al*., 2015) on AGO2 immunoprecipitates when we transfected control or QKI shRNA in HeLa cells. Similar to the enrichment of let-7b on AGO2, the depletion of QKI decreased enrichment of *POLR2D* mRNA on AGO2 (**Figure 4A *Left*)**. The QKI effects on interaction of AGO2 and *POLR2D* mRNA inversely correlate on *POLR2D* mRNA steady-state levels (**Figure 4A *Right***), possibly due to changes of *POLR2D* mRNA stability but not changes in AGO2 level (**Figure 4B**). To measure the stability of *POLR2D* mRNA after knocking down QKI (or luciferase as a control), *POLR2D* and *GAPDH* mRNA were measured at select times (0, 2, 4, 6, and 8 hours) after treatment with Actinomycin D. We observed that the stability of *POLR2D* mRNA increases as the half-life of *POLR2D* mRNA (t_1/2,_ 50% of original abundance) increases from 4.2 hours to 7.8 hours after silencing of QKI in HeLa cells (**Figure 4C**). In contrast, the stability of *GAPDH* mRNA, (the house-keeping mRNA), does not change.

**Figure 4.**
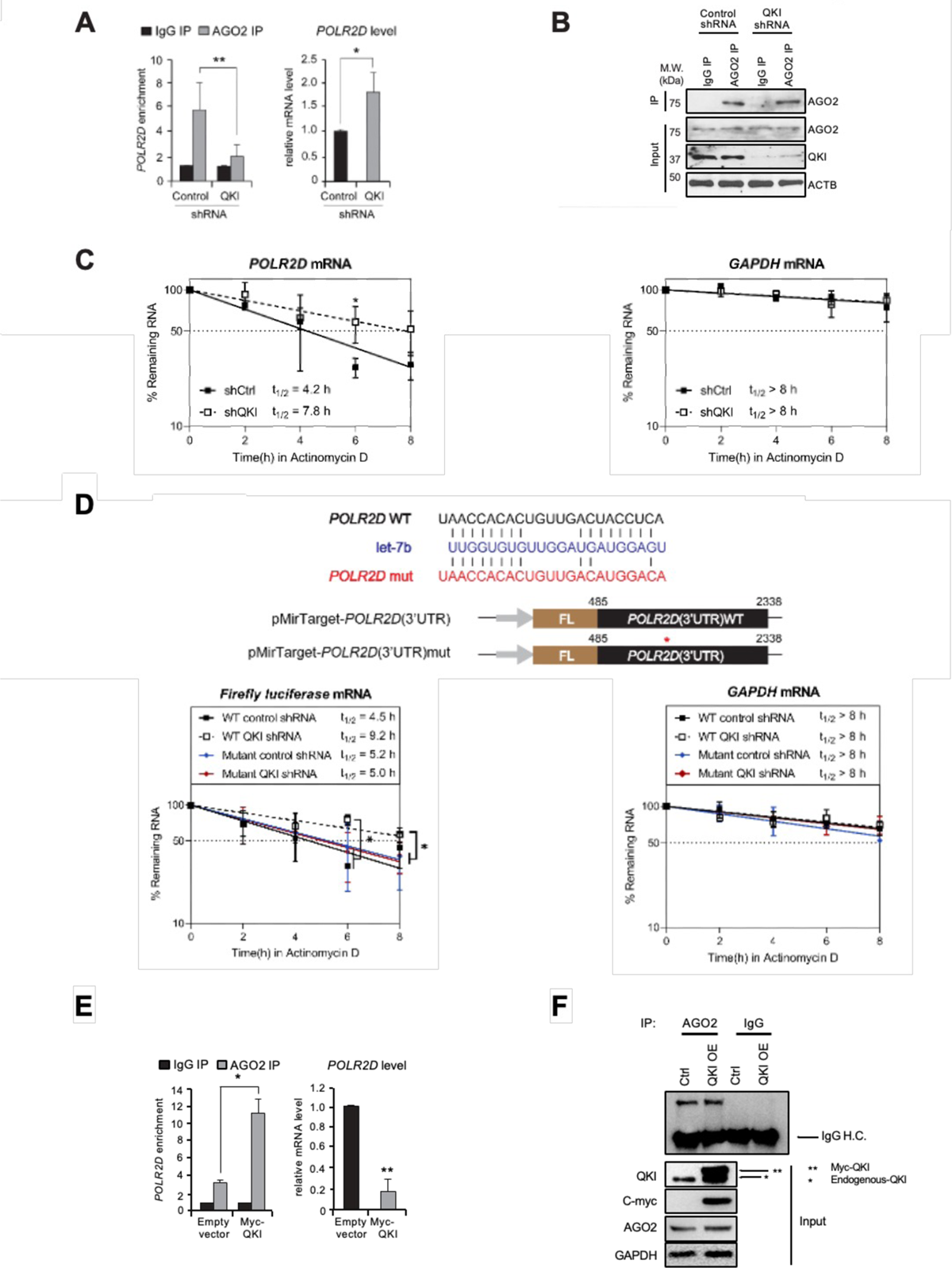
QKI is essential for let-7b-mediated mRNA decay. **(A)** AGO2 RIP assay was performed in HeLa cells after transfection of control and QKI shRNA for detection of *POLR2D* mRNA. (*Left*) After AGO2 RIP, the amount of *POLR2D* mRNA (a let-7b target mRNA) from immune pellets were determined by reverse transcription (RT) and qPCR, with normalization to *TUBA* mRNA. (*Right*) Total RNAs were extracted from control and QKI shRNA cells for RT-qPCR. N = 3, \*\**p* < 0.01, \**p* < 0.05, not significant from Student’s *t*-test. Data are mean ± S.D. **(B)** AGO2 RIP samples were subjected to SDS-PAGE for western blot analysis against AGO2, QKI and ACTB. **(C)** Measurement of *POLR2D* and *GAPDH* mRNA stability with Actinomycin D treatment after transfection of control or QKI shRNA in HeLa cells. At a given time of Actinomycin D treatment, total RNA was purified and converted into cDNA for quantitation of *POLR2D*, *GAPDH* mRNA, or *18S* RNA. The stability of (*Left*) *POLR2D* and (*Right*) *GAPDH* mRNAs were determined by comparing the remaining amounts of mRNA to the pre-treatment levels, which were normalized using 18S RNA. N = 3, \**p* < 0.05, *n.s.*, not significant from Student’s *t*-test. Data are mean ± S.D. **(D)** Measurement of *LUC2* reporter and *GAPDH* mRNA stability after transfection of control and QKI shRNA in HeLa cells. After treatment of Actinomycin D, stability of *LUC2 (Left)* and *GAPDH* mRNAs (*Right*) were assessed by RT-qPCR with normalization of *18S* RNA (WT QKI shRNA vs WT control/mutant control/mutant QKI). N = 3, \**p* < 0.05, *n.s.*, not significant from Student’s *t*-test. Data are mean ± S.D. **(E)** AGO2 RIP assay using lysates from HeLa cells after transfection of empty vector and Myc-QKI overexpression plasmids for detection of *POLR2D* mRNA. (*Left*) After AGO2 RIP, the amount of *POLR2D* mRNA from immune pellets were determined by RT-qPCR with normalization to *TUBA* mRNA. (*Right*) Steady-state level of *POLR2D* mRNA was measured by RT-qPCR using total RNAs extracted from control and QKI shRNA. N = 3, \*\**p* < 0.01, \**p* < 0.05, from Student’s *t*-test. Data are mean ± S.D. **(F)** AGO2 IP efficiency was verified after AGO2 immunoprecipitation and western blot analysis using lysates from HeLa cells transfected with control or Myc-QKI plasmids.

We further compared the stability of luciferase reporter mRNA containing the 3’ UTR of *POLR2D* mRNA (Yoon *et al*., 2015) in HeLa cells transfected with control or QKI shRNA. While the half-life of the Firefly luciferase reporter mRNA increases from 4.5 hours to 9.2 hours with transfection of QKI shRNA, the half-life of mutant reporter mRNA lacking the let-7b seed-sequence was resistant to lowering QKI levels (t_1/2_ = 5.0 hours) (**Figure 4D**). As an additional control, we examined *GAPDH* mRNA, the stability of which did not change significantly. Overexpression of QKI increased the enrichment of *POLR2D* mRNA on AGO2 (**Figure 4E *Left***) and decreased steady state level of *POLR2D* mRNA (**Figure 4E *Right***) without changes of AGO2 level or immunoprecipitation efficiency (**Figure 4F**). To further investigate QKI’s role in miRNA-mediated gene silencing, we generated a luciferase reporter with *cMYC* 3’ UTR spanning let-7b binding sites as predicted in mirTarBase database to confirm the influence of QKI on let-7b activity. We cloned the full-length and minimum containing let-7b binding sites of *cMYC* 3’ UTR into a dual luciferase reporter construct and then co-transfected these plasmids with either QKI or control plasmid into HeLa cells. QKI overexpression indeed decreased the luciferase activity which is regulated by *cMYC* 3’ UTR, indicating that QKI promotes let-7b-mediated gene silencing (**Figure S4A**). We also observed downregulation of cMYC expression by western blot analysis after QKI overexpression (**Figure S4B**). cMYC is associated with tumor progression involving in cell proliferation and migration (Dang, 2012; Meškytė *et al*, 2020). Given that QKI suppresses cMYC expression, we further investigated the impact of QKI on cell proliferation and migration. Overexpression of QKI decreased proliferation and migration of HeLa cells (**Figure S4C and S4D**). These results demonstrate that QKI suppresses cell proliferation and migration by enhancing let-7b-mediated gene silencing.

A previous study reported that QKI is immunoprecipitated with AGO2 and C-terminal domain of QKI is responsible for interaction with AGO2 in human cell lines (Wang *et al*, 2010a). To test whether increased let-7b activity by QKI was due to the interaction between AGO2 and C-terminal domain of QKI, we constructed Myc-tagged C-terminal truncated QKI (**Figure 5A, *Left***) and performed immunoprecpitation using anti-Myc or anti-AGO2 antibodies followed by RT-qPCR in QKI-depleted cells. Our data revealed that exogenous QKI can be associated with let-7b, but not QKI mutant lacking C-terminal domain, suggesting that C-terminal domain may be required for interaction with let-7b (**Figure 5A, *Middle***). AGO2 RIP also showed the restoration of let-7b enrichment on AGO2 immunoprecipitates when wild-type QKI is reintroduced compared to C-terminal mutant QKI (**Figure 5A, *Right***). We confirmed steady state levels of let-7b were not changed significantly suggesting that QKI may neither influence miRNA biogenesis nor miRNA stability (**Figure 5B**). Consistently, increased of *POLR2D* mRNA stability by QKI depletion was decreased by re-introduction of wild type QKI but not by QKI-C-terminal truncated mutant (**Figure 5C)**. The results demonstrate that interaction of QKI with AGO2 could be important for transferring miRNAs which subsequently enhances target RNA decay. Additionally, overexpression of AGO2 in QKI-depleted cells decreased stability of *POLR2D* mRNA (**Figure 5D**) without affecting level of QKI proteins (**Figure 5E**), indicating that QKI-mediated *POLR2D* mRNA decay is dependent on miRNA function. These results demonstrate that QKI promotes decay of let-7b target mRNAs indirectly by transferring let-7b to AGO2. The latter action may be achieved through multiple mechanisms: decapping, deadenylation, or endonucleolytic cleavage, and further studies are needed to fully confirm and specify their role in the degradation.

**Figure 5.**
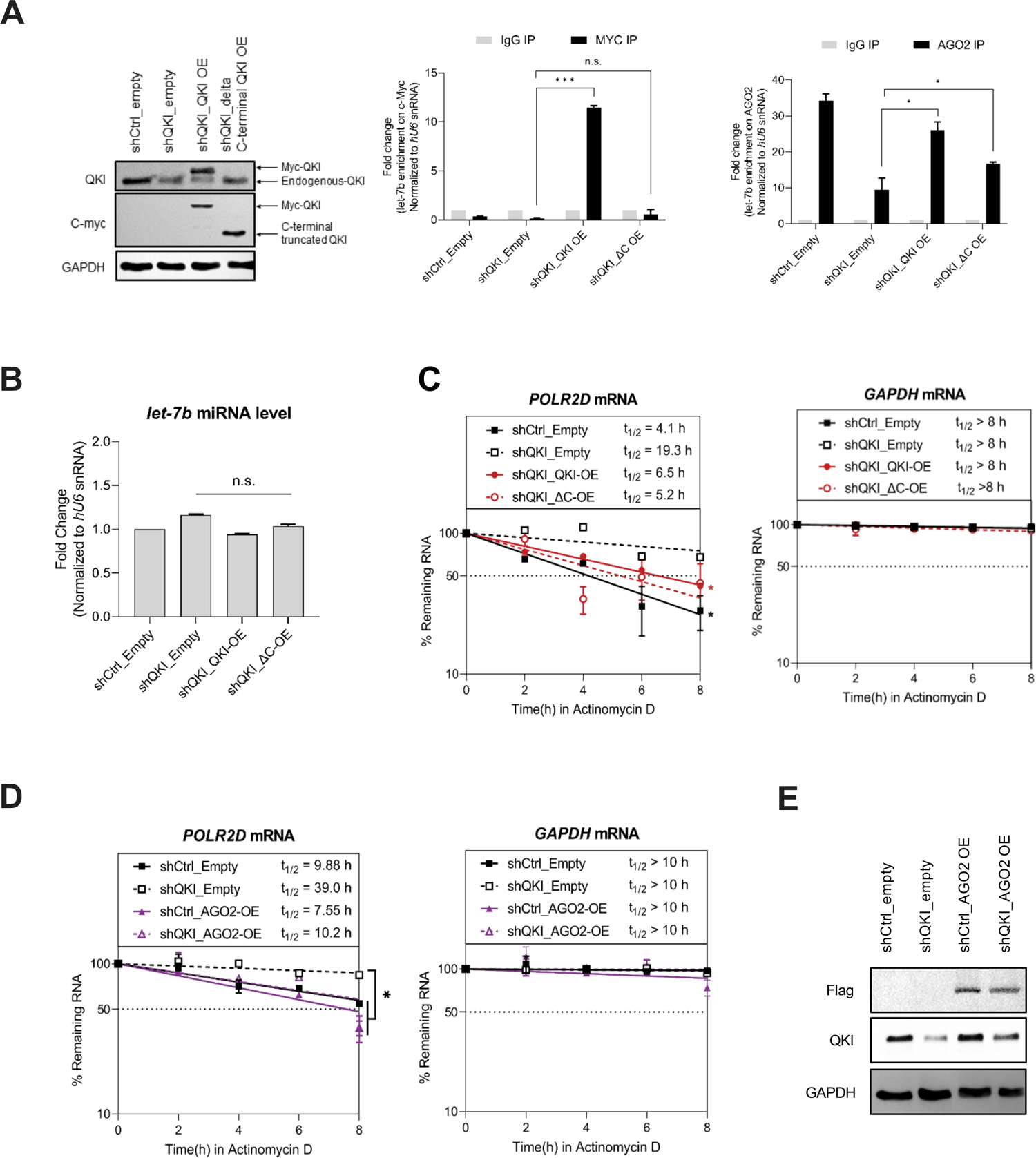
C-terminal region of QKI is essential for AGO2-mediated mRNA decay. **(A)** (*Left*) AGO2 RIP samples were subjected to SDS-PAGE for western blot analysis against QKI, c-Myc and GAPDH. (*Middle*) MYC RIP and (*Right*) AGO2 RIP and let-7b RT-qPCR assay were performed in HeLa cells transfected with control shRNA or QKI shRNA in combination with QKI full length or CK mutant plasmids. At 48 hours after transfection, cell lysates were subjected to immunoprecipitation against AGO2 for enrichment of co-purified let-7b by RT-qPCR. N = 2, \*\*\**p* < 0.001, \**p* < 0.05, *n.s.*, not significant from Student’s *t*-test. Data are mean ± S.D. **(B)** RT-qPCR levels of let-7b using RNAs purified from the corresponding samples. N = 2, *n.s.*, not significant from Student’s *t*-test. Data are mean ± S.D. **(C)** Measurement of *POLR2D* mRNA (*Left*) and *GAPDH* mRNA stability (*Right*) after re-introducing QKI full length or CK mutant in QKI knockdown cells. After treatment of Actinomycin D, stability of *POLR2D* and *GAPDH* mRNAs were assessed by RT-qPCR with normalization of *18S* RNA. N = 2, \**p* < 0.05, *n.s.* not significant from Student’s *t*-test. Data are mean ± S.D. **(D)** Measurement of *POLD2D* mRNA (*Left*) and *GAPDH* mRNA (*Right*) stability after transfection of Flag-tagged AGO2 expression or empty vector in HeLa cells either stably expressing control shRNA or QKI shRNA. After treatment of Actinomycin D, stability of *POLR2D* and *GAPDH* mRNAs were assessed by RT-qPCR with normalization of *18S* RNA. N = 2, \**p* < 0.05, *n.s.*, not significant from Student’s *t*-test. Data are mean ± S.D. **(E)** *POLR2D* mRNA stability assay samples were subjected to SDS-PAGE for western blot analysis against Flag, QKI and GAPDH antibodies.

### QKI suppresses assembly of AGO2-let-7b-target mRNA complex, and dissociation of let-7b from AGO2

Possible mechanisms by which QKI promotes target mRNA decay are to accelerate assembly of the AGO2-let-7b-target mRNA complex, and to inhibit release of miRNA from AGO2. We next performed *in vitro* miRNA release assay whether QKI can repress dissociation of let-7b from AGO2 (Min *et al*, 2020). We incubated recombinant AGO2 (20 nM) with let-7b (20 pM), followed by immunoprecipitation of AGO2, and the removal of free AGO2 after several washing steps. We then compared the amount of let-7b from the supernatant after incubation of AGO2-let7b complex, in the presence and absence of recombinant QKI (**Figure 6A**). Our RT-qPCR of let-7b revealed that the addition of recombinant QKI significantly suppresses let-7b dissociation from AGO2. Similarly, we observed suppression of let-7b and *POLR2D* mRNA release from AGO2 immunopellets by recombinant QKI using HeLa cell lysates (**Figure 6B**). These results demonstrate that QKI maintains the interaction of AGO2 and let-7b. In order to further substantiate our hypothesis which is that QKI promotes let-7b mediated target gene silencing, we performed proximity ligation followed by PCR using HeLa cell lysates (Broughton *et al*, 2016). In this assay, we crosslinked AGO2/let-7b/target RNA complexes using UV-C light, digested protein-unbound RNA for internal ligating let-7b with target RNA fragment, and then immunoprecipitated AGO2. After immunoprecipitation, intermolecular ligation step was performed to ligate let-7b with their target RNAs. Let-7b-*POLR2D* mRNA chimeric RNA fragment were subsequently converted to cDNA, and then relative amount were compared between control and QKI-depleted HeLa cells **(Figure S5A**). As expected, we observed that depletion of QKI decreases the amount of let-7b-*POLR2D* chimeric product bound to AGO2, indicating that QKI stabilizes let-7b and *POLR2D* mRNA interaction (**Figure S5B**). Moreover, we also profiled total RNA and focused on changes in let-7b target RNAs from control and QKI-depleted HeLa cells. The result showed that let-7b target RNAs are prone to more presence when QKI is depleted (**Figure S5C**). These findings show that QKI inhibits release of let-7b, and targets mRNA from the AGO2-containing gene-silencing complex.

**Figure 6.**
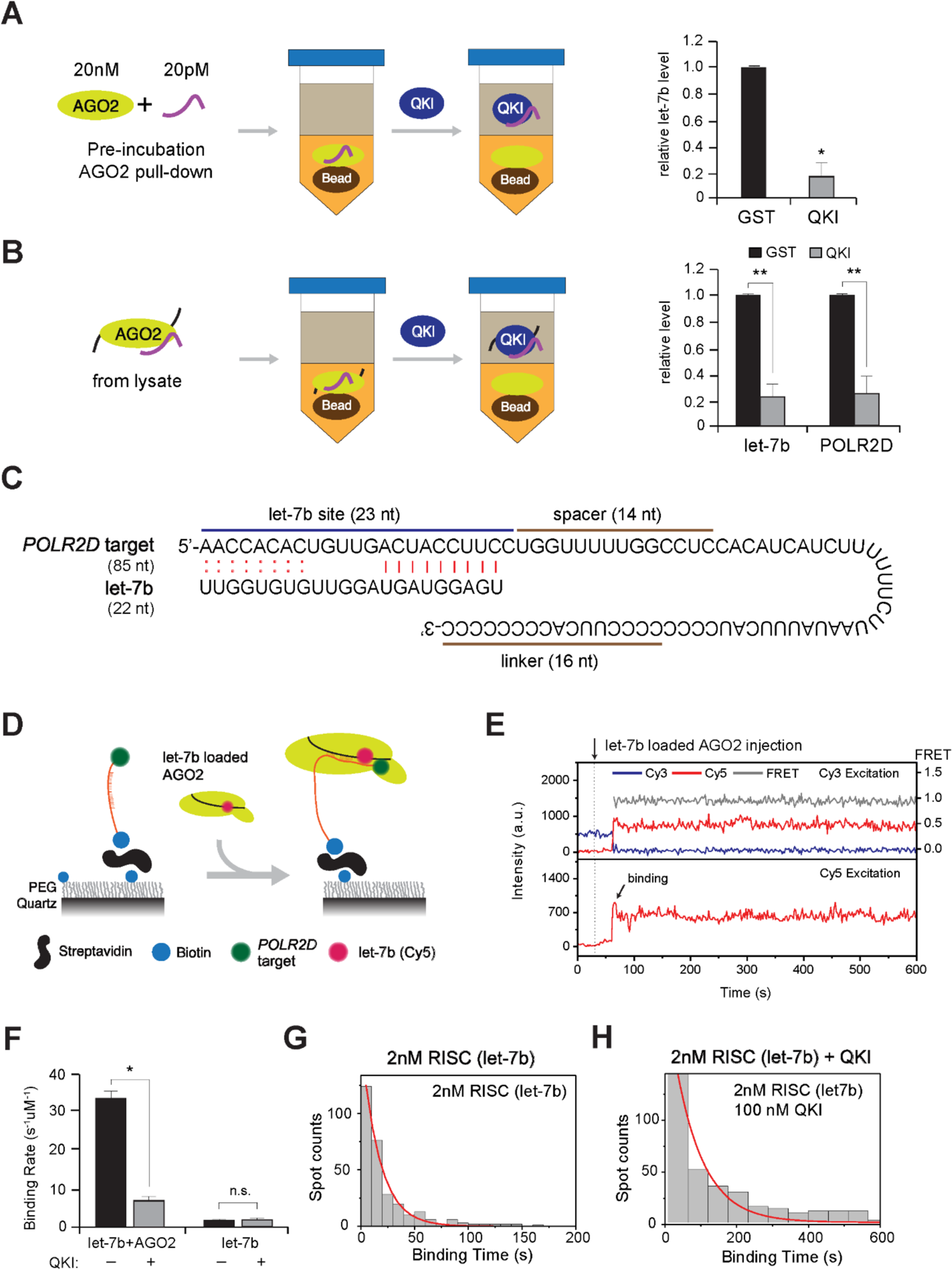
QKI slows down AGO2-let-7b-target RNA assembly and suppresses let-7b release. **(A)** Detection of let-7b released from recombinant AGO2 *in vitro*. AGO2 in complex with let-7b was immobilized on the sepharose beads for detection of let-7b from the supernatant, in the presence of GST or QKI using RT-qPCR. N = 3, \**p* < 0.05, from Student’s *t*-test. Data are mean ± S.D. **(B)** Detection of let-7b and *POLR2D* mRNA released from AGO2 *in vivo*. AGO2 in complex with let-7b and *POLR2D* mRNA was immobilized on the sepharose beads using HeLa cell lysates for detection of let-7b from the supernatant, in the presence of GST or QKI using RT-qPCR. N = 3, \*\**p* < 0.01, from Student’s *t*-test. Data are mean ± S.D. **(C)** Target RNA sequence of *POLR2D* 3’ UTR fragment containing let-7b site. Cy3 was labeled at the 5’-end and biotin was labeled at the 3’-end. **(D)** Single-molecule assay of RISC assembly with target RNA in the presence of QKI. The RISC solution (AGO2 and let-7b) was injected into the chamber in the presence or absence of excessive QKI (100 nM) with pre-incubation for 5 minutes. **(E)** Representative time trace of fluorescence of Cy3 on *POLR2D* fragment. The RISC solution was injected into the chamber at 40 s (dashed line). **(F)** The binding rate of let-7b to *POLR2D* target was obtained through single exponential fit of the binding time histogram. The number of data is > 330 each. N = 3, \**p* < 0.05, *n.s.*, not significant from Student’s *t*-test. Data are mean ± S.D. **(G, H)** Histograms of binding time in (G) absence or (H) presence of recombinant OKI. The red lines are single exponential decay fit (17 s and 70 s, respectively), N = 3.

Next, we tested if QKI affects assembly of AGO2-let-7b-target mRNA complex at the single-molecule level, as performed previously (Jo *et al*, 2015; Min *et al*, 2017). We immobilized the fragment of *POLR2D* 3’ UTR containing the let-7b target sequence (**Figure 6D**), through a streptavidin–biotin interaction. The 5’ end of the target RNA was labeled with Cy3 to induce high FRET (Fluorescence Resonance Energy Transfer) when Cy5-labeled let-7b is bound on the target RNA. The Cy5-labeled let-7b was pre-incubated with an excess of AGO2 (25X, 1 μM), and the diluted reaction mixture (2 nM) was injected into the detection chamber (**Figure 6E**). In this setting, the simultaneous appearance of high FRET for Cy3 excitation, and the resulting Cy5 signal, indicates the binding of the RISC on target RNA. The binding rate was measured by collecting the time delay between the solution injection and high FRET signal from the binding events. Our results revealed that the assembly of AGO2-let-7b-*POLR2D* fragment complex significantly slowed down about five times when recombinant QKI was added (100 nM) over the indicated times (**Figure 6F-H**). The *in vitro* kinetic assay showed that QKI significantly slows down the assembly of this complex, and suggest that QKI suppresses the kinetics of AGO2-let-7b-target mRNA assembly.

## DISCUSSION

MiRNAs and RBPs, including miRBPs, are key regulators of post-transcriptional gene expression (Ciafrè & Galardi, 2013; Jonas & Izaurralde, 2015; Min *et al*., 2017). Dysregulation of miRNA target recognition, miRNA biogenesis, and their subcellular localization, alters cellular physiology, and drives human pathologies such as cancer, inflammation, cardiac abnormalities, ischemia/stroke, and neuronal degeneration (Abe & Bonini, 2013; Chen *et al*, 2017; Ding *et al*, 2011; Rupaimoole *et al*, 2016; Zangari *et al*, 2017). Thus, investigation of interplay between miRNAs and miRBPs will contribute to find clues to treat human diseases by targeting either miRNAs or miRBPs.

Although QKI has been studied by splicing of precursor mRNA in nucleus (Wu *et al*, 2002), none of the previous studies reported QKI as a mature miRNA-binding protein. Our new findings show that QKI binds directly to let-7b miRNA, and accelerates AGO2/let-7b*-*mediated target mRNA decay (**Figure 1 and 4**). We also revealed that QKI acts as a tumor suppressor gene by partially increasing the activity of let-7b in HeLa cells (**Figure S4**). QKI may empower miRNA-mediated gene silencing when QKI expression or cytoplasmic localization changes. We do not know the exact molecular mechanisms by which QKI decreases the stability of let-7b target mRNA, because QKI transfer of let-7b to AGO2: (i) is inefficient (**Figure 3**), (ii) suppresses the kinetic assembly of AGO2/let-7b/*POLR2D* mRNA at the single-molecule level, and (iii) inhibits disassembly of AGO2/let-7b/*POLR2D* mRNA complex (**Figure 6**). Even though we confirmed through chimera PCR in cell lysates that QKI is critical for let-7b and *POLR2D* mRNA interaction (**Figure S5**), the detailed mechanistic function by which QKI stabilizes AGO2/let-7b/target RNA remains to be determined.

### Presence of mature miRNA without RISC assembly and novel miRBPs in addition to AGO-family proteins

Mature miRNAs function in mRNA decay and translation as a part of RISC (Misiewicz-Krzeminska *et al*, 2019). It has been reported that miRNAs dissociated from AGO are rapidly degraded by mechanisms of Target-Directed MicroRNA Degradation (TDMD) (Han *et al*., 2020; Kingston & Bartel, 2021; Kleaveland *et al*., 2018; Shi *et al*., 2020). Previous studies also revealed that levels of a subset of mature miRNAs correlate with levels of Argonaute proteins (Diederichs *et al*, 2008; Martinez & Gregory, 2013; O’Carroll *et al*, 2007; Reichholf *et al*., 2019). However, our data showed that the presence of AGO-free mature miRNAs existed in cytoplasm, which is supported by using RNA-seq of AGO TKO cells data and AGO IP data. It indicated that there are other RBPs binding with mature miRNA. Certain RBPs interact with mature miRNAs directly at high affinity (Zealy *et al*, 2017); however, it is not known if these miRBPs also interact with duplex miRNAs. The results of our PAR-CLIP analysis revealed a subset of mature miRNA as QKI targets (**Figure S1D**), not passenger strands. Differences in amino acid sequences and structures between QKI and AGO proteins also suggest that these two miRBPs have different modes of miRNA interactions, such as recognition of 5’ phosphoryl and 3’ hydroxyl groups. It is also possible that miRBPs modulate miRNA-mediated gene silencing by functioning as auxiliary factors surveilling target recognition by AGO proteins (Min *et al*., 2017; Yoon *et al*, 2013b), or simply compete with AGO proteins for miRNAs (Eiring *et al*, 2010). *k*_a_ of recombinant QKI for let-7b (3.24 x 10^4^ M^-1^s^-1^) slower than a wide range of eukaryotic proteins (Gleitsman *et al*, 2017) but several proteins indeed exhibit *k_on_* in a similar range of *k_a_* for QKI with let-7b (Arluison *et al*, 1999; Chen *et al*, 2005; Park-Lee *et al*, 2003; Park *et al*, 2000; Salomon *et al*, 2015; Shen & Shan, 2010). Although we cannot exclude the possibility that QKI may not facilitate the direct loading of let-7b to AGO2, other components would be necessary to facilitate the direct loading of let-7b to QKI, or that the transfer is not very efficient. Further studies are needed to identify common sequences and, or structural motifs among miRBPs to allow their interaction with mature miRNAs such as let-7b.

### Mechanisms of miRNA-mediated gene silencing via delay of RISC assembly and disassembly

The discrepancy between results of cell-based biochemistry and single-molecule biophysics suggests that there should be additional factors contributing to efficient RISC assembly, such as scaffolding proteins supporting QKI and AGO2 interaction in specific locations (Wang *et al*., 2010a). A recent study argues that the AGO2 phosphorylation cycle regulates the interaction of AGO2 with target mRNAs, to maintain the global efficiency of miRNA-mediated gene silencing (Golden *et al*, 2017). The former study (Golden *et al*., 2017) identified a protein kinase (CSNK1A1) and a phosphatase complex (ANKRD52-PPP6C) involved in the hierarchical cycle of AGO2 phosphorylation. AGO2 phosphorylation could induce active disassembly of AGO2 and target mRNA complexes, giving rise to increase availability of AGO to silence additional targets. Considering this, the kinetic repression of AGO2-miRNA-target mRNA assembly by QKI *in vitro* (**Figure 6**) could promote mRNA decay in mammalian cells. Future studies could investigate whether QKI influences AGO2 phosphorylation by interacting with the aforementioned kinase and phosphatase complex, and/or regulating target mRNA silencing.

In summary, our findings add growing evidence that novel miRBPs occupy AGO2-free mature miRNAs functioning in miRNA metabolism. Our observations emphasize the regulation of AGO2/miRNA/target mRNA assembly by QKI, which can impact upon target gene silencing. Our study provides an insight upon how miRNA-binding activity of QKI is linked to cellular processes impacting miRNA metabolism.

## METERIALS and METHODS

### Contact for Reagent and Resource Sharing

Further information and requests for resources and reagents should be directed to and will be fulfilled by the Lead Contact, Je-Hyun Yoon.

### Experimental Models and Subject Details

#### Cell culture, transfection, short hairpin RNAs, and plasmids

Human HeLa, Mouse Embryonic Fibroblast (MEF), and HCT116 cells were cultured in DMEM (Invitrogen) supplemented with 10% (v/v) FBS and antibiotics. Using PEI (polyethylenimine), the cells were then transfected with shRNA plasmids designed to luciferase (5’-CGCTGAGTACTTCGAAATGTC-3’), or QKI (5’-GCTCAGAACAGAGCAGAAATC-3’). pcDNA3.1 and pDESTmycQKI (Addgene, #19870) were used for overexpression experiments. Cells were analyzed for RT-qPCR, western blot, and exosome purification 48 hours after transfection. To construct C-terminal deletion mutant (CK-truncated) QKI, the corresponding DNA region was first amplified from the pDESTmycQKI. The resulting PCR products were then sub-cloned the the *SalI* and *NotI* sites of the pDESTmycQKI plasmid after removing full length QKI sequences. To construct luciferase reporter vector containing the 3’UTR of *cMYC* mRNA, the full-length (FL) and minimum (Min) containing let-7b binding site of 3’ UTR of *cMYC* was cloned into the pmiRGLO Dual-Luciferase miRNA target expression vector (Promega). *cMYC* 3’ UTR region was first amplified from genomic DNA. Primer sequences are listed in Table S6.

#### RNP analysis

For immunoprecipitation (IP) of RNP complexes from cell lysates [RIP analysis] (Yoon *et al*, 2012), cells were lysed in Protein Extraction Buffer (PEB) with 20 mM Tris-HCl at pH 7.5, 100 mM KCl, 5 mM MgCl_2_ and 0.5% NP-40 for 10 min on ice, and centrifuged at 10,000 × g for 15 min at 4°C. The supernatant was then incubated 1 hour at 4°C with protein A-Sepharose beads coated with antibodies recognizing QKI (ab126742), AGO2 (ab57113), Myc (sc-40), or control IgG (Santa Cruz Biotechnology). After the beads were washed with NT2 buffer (50 mM Tris-HCl at pH 7.5, 150 mM NaCl, 1 mM MgCl_2_ and 0.05% NP-40), the complexes were incubated for 15 minutes at 37°C with 20 units of RNase-free DNase I. They were finally incubated for 15 minutes at 55 °C with 0.1% SDS and 0.5 mg/ml Proteinase K, to remove remaining DNA and proteins, respectively. The RNA isolated from the IP materials was further assessed by Reverse transcription and quantitative PCR (RT-qPCR) analysis using the primers (**Table S6**). Normalization of RIP results was carried out by quantifying, in parallel, the relative levels of housekeeping RNAs such as *GAPDH* mRNA and *U6* snRNA in each IP sample. These abundant RNAs are non-specifically recovered during IP reactions.

#### Western blot analysis

Whole-cell lysates prepared in PEB were separated by SDS-polyacrylamide gel electrophoresis (SDS-PAGE) and transferred onto nitrocellulose membranes (Invitrogen iBlot Stack). Anti-β-Actin (sc-47778) and anti-cMyc (sc-40) antibodies were purchased from Santa Cruz Biotechnology. Anti-Dicer (ab82539), anti-AGO2 (ab57113) and anti-QKI (ab126742) antibodies were purchased from Abcam. anti-GAPDH (2118S), endogenous c-Myc (D84C12) antibodies were purchased from Cell Signaling Technology. Anti-pan AGO (MABE56) and anti-Flag antibody (SLBX2256) antibodies were purchased from Sigma. The HRP-conjugated secondary antibodies were purchased from GE Healthcare.

#### RNA analysis

For RIP analysis, Ribozole (Amresco) was used to extract total RNA, and acidic phenol (Ambion) was used to extract RNA. Reverse transcription (RT) was performed using random hexamers and reverse transcriptase (Maxima, Thermo Scientific), and real-time, quantitative (q)PCR using gene-specific primers (**Table S6**), and SYBR green master mix (Kapa Biosystems), using a BioRad iCycler instrument.

MicroRNA quantitation was performed after RNA extraction from immunoprecipitated samples (with anti-AGO2 antibody or control IgG), polyadenylation (QuantiMiR kit), and hybridization with oligo-dT adaptors. After RT, cDNAs were quantitated by qPCR with microRNA-specific primers, or with primers to detect the control transcript *U6* snRNA, along with a universal primer. *U6* snRNA was used a normalization of exosomal miRNA (Xu *et al*, 2017; Zhang *et al*, 2018)

#### Single-molecule assay

This assay was performed according to modifications of a previously described method (Jo *et al*., 2015) (Yoon *et al*., 2015). Quartz slides and coverslips were cleaned in piranha solution (3:1 solution of concentrated sulfuric acid and 30% [v/v] hydrogen peroxide) for 20 min, and coated with 40:1 mixture of polyethylene glycol (MPEG-SVA-5000; Laysan Bio) and biotinylated polyethylene glycol (Biotin-PEG-SVA-5000; Laysan Bio). A detection chamber was assembled by sandwiching double-sided sticky tape (3M) between a quartz slide and a coverslip. Polyethylene tubing (PE50; Becton Dickinson) was connected to the inlet and outlet of the chamber for stable buffer exchange during fluorescence signal measurement. For surface immobilization of target RNAs, streptavidin (0.2 mg/ml; Invitrogen) and target RNA (50 pM) were sequentially injected into the flow-cell (detection chamber) and incubated for 2 min. To establish the AGO2:let-7b complex, let-7b miRNA (40 nM) was incubated with excess human Argonaute 2 (1 μM; Sino Biological) at 23°C for 1 hour.

Fluorescence measurements were carried out with an oxygen-scavenging imaging buffer which contained 20 mM Tris-HCl (pH 8.0), 135 mM KCl, 1mM MgCl_2_, 1 mM Trolox (Sigma), 1 mg/mL glucose oxidase (Sigma), 0.04 mg/ml catalase (Sigma), 0.4% (w/v) glucose (Sigma), and 2000 U/ml RNase inhibitor (Promega). A single-molecule fluorescence image was taken using a home-built prism-type Total Internal Reflection Fluorescence Microscope (TIRFM) equipped with an electron-multiplying CCD camera (IXON DV597ECS-BV; Andor Technology) at a frame rate of 1 Hz (Roy *et al*, 2008) with alternating laser excitation. The temperature of the flow-cell and solution was maintained at 37°C with a temperature control system (Live Cell Instruments).

#### High throughput RNA sequencing

Total RNA was isolated from human HeLa cells transfected with control or QKI shRNA using TRIzol according to manufacturer’s instructions. For RNA-sequencing, we followed standard Illumina sequencing protocols using the TruSeq Stranded Total RNA kit and NovaSeq6000 S4 (150bp PE). The resulting data were aligned to human genome sequences (HG19) for assembly and quantitation.

#### Small RNA sequencing and analysis

Library preparation was performed using the TruSeq Small RNA library preparation kit (Illumina, RS-200) following the manufacturer’s instructions. Briefly, 1 μg of input RNA was loaded into Urea-TBE gels for purification and the resulting RNAs were applied for the following procedures. User-supplied reagents including T4 RNA ligase2 Deletion Mutant (Lucigen, LR2D1132K) and Maxima First Stand cDNA synthesis kit (Thermo Fisher Scientific, K1641) were purchased separately. Libraries were amplified using 11 cycles of PCR for the manufacture’s index or modified index primer set to increase diversity. Libraries were prepared according to the manufacturer’s protocol with a modification in the size selection step, which, instead of agarose gel purification, PippinHT Prep instrument (Sage Science, HTP0001) and 3% agarose dye-free cassette with internal standards (Sage Science, HTG3010) was used under the following conditions: base pair start = 120 bp, base pair end = 160 bp, range = broad, target peak size = 145 bp. Eluted Libraries from PippinHT system were subsequently analyzed on Tape-station 4150 (Agilent Technologies, G2992AA) following the manufacturer’s instructions using a High Sensitivity DNA Screen tape (Agilent Technologies, 5067-5584). Each library was barcoded with unique sequence of reverse primers during the PCR step, which contained Illumina compatible indices and modified indexes (see GEO database). Before pooling libraries for the next-generation sequencing, concentration of each library was measured in a high sensitivity Tape-station followed by smear analysis. Illumina NextSeq 550 or Miniseq with single end (50 nt;R1) read method was apply for the library sequencing.

Quality control and trimming of sequencing reads were performed using fastp (ref; fastp: an ultra-fast all-in-one FASTQ preprocessor) with default parameter setting. Subsequently, reads shorter than 15 bases were additionally excluded for the downstream analysis. The trimmed reads were mapped based on mature and hairpin databases (Kozomara *et al*, 2019) for respective species (mouse or human) after further clipped 3’ Illumina stop oligo sequence (GAATTCCACCACGTTCCCGTGG). Then they were quantified, and RPM (reads per million mapped reads) normalized. The mapping and quantification steps were performed by mirDeep2 (Friedländer *et al*, 2012).

#### Chimera PCR

This assay was slightly modified from previously described method (Broughton & Pasquinelli, 2018). In brief, cells were crosslinked using UV-C (254nm) crosslinker. After UV crosslinking, cells were lysed with lysis buffer (20mM Tris-HCl pH 7.4, 10mM KCl, 5mM MgCl_2_, 0.5% NP-40 supplemented with protease inhibitor cocktail, DTT, and RNase inhibitor), sonicated for 10 sec and centrifuged at 17,000 x g for 15 min. Lysates were treated with RNase I (NEB) and Turbo DNase (Invitrogen) to digest RNA not protected by Argonaute. Lysates were then immunoprecipitated using anti-AGO2 antibody (Abcam) and Protein G dynabeads (Invitrogen). Beads were washed with high-salt wash buffer (50mM Tris-HCl pH 7.4, 1M NaCl, 1mM EDTA, 1% NP-40, 0.1% SDS, 0.5% sodium deoxycholate) and low-salt wash buffer (20mM Tris-HCl pH 7.4, 10mM MgCl_2_, 0.2% Tween-20). The 5’ end of target RNA was phosphorylated using T4 PNK 3’ phosphatase minus (NEB) and 1mM ATP (NEB) for 10 min at 37°C with shaking. The beads were then washed with low-salt wash buffer. Intermolecular ligation of miRNA and target RNA was performed with T4 RNA ligase I (NEB), 1mM ATP and PEG400 for overnight at 16°C. The beads were then washed with low-salt wash buffer. The 3’ phosphate left over after RNase I digestion was removed using T4 PNK (NEB) and the ligation of the 3’ linker was performed using T4 RNA ligase Ⅰ (NEB). After linker ligation, the RNA was isolated using proteinase K and reverse transcribed. cDNAs were then size selected to remove extra reverse transcription primer using AMPure XP beads (Beckman Coulter). The cDNAs were circularized using CirLligase Ⅱ (Lucigen), linearized using BamHⅠ (NEB) and libraries were then amplified using Taq DNA polymerase (Enzynomics). Libraries containing residual primers were purified by AMPure XP beads before specific-chimera amplification. Let-7b-*POLR2D* chimeras were detected by PCR amplification using a forward primer matching the mature let-7b sequence and a reverse primer complementary to the *POLR2D* target site and were detected by DNA polyacrylamide gel electrophoresis. Primer sequences are listed in Table S6.

#### miRNA activity reporter assay

HeLa cells were seeded at a density of ∼60% in a 12-well culture plate and incubated at 37°C. After 24 hours of transfection, the cells were washed twice with PBS. Dual luciferase assay was performed using the Dual luciferase reporter assay system (Promega, #E1910) according to manufacturer’s instructions.

#### Migration assay

Cell migration assays were performed using Transwell plates with 8.0 μm pore size membranes (Falcon, #353097). The cells were seeded at 2.5 x 10^5^ in 200 μl of serum-free medium and added to the upper chamber of a 24-well plate. The lower chamber was filled with 750 μl of medium with serum. The 24-well plate was incubated at 37°C for 24 hours. Migrated cells on the bottom side of the membranes were fixed with 4% formaldehyde in PBS for 15 minutes and the chamber was then washed with PBS. The cells were permeabilized with PBS containing 0.25% X-100 for 10 minutes and the chamber was washed again with PBS. Cells on the lower chamber membrane were stained with 0.5% crystal violet dissolved in 20% methanol for 15 minutes and then the chamber was washed with PBS. Non-migrating cells were scraped off with cotton swabs. Migrated cells were quantified using ImageJ.

#### Cell Proliferation assay

The cell proliferation was measured using the EZ-CYTOX (DoGen, #EZ-3000,) according to manufacturer’s instructions. Briefly, cells were seeded at 4 x 10^3^ cells/well in 100 µl of culture medium on a 96-well plate. After 24 hours, the cells were washed with PBS. The culture medium and EZ-CYTOX reagent were then diluted at a ratio of 9:1 and 100 µl of the mixture was added to each well. The 96-well plate was incubated at 37°C for 30 minutes. The absorbance at 450 nm was then measured using a plate reader.

## ACKNOWLEDGEMENTS

We thank D. Patel (Memorial Sloan Kettering Cancer center, NY, USA) and MJ Gamble (Albert Einstein College of Medicine, NY, USA) for providing plasmids. We appreciate GM Wilson (University of Maryland School of Medicine, MD, USA) for providing critical comments.

## DECLARATION OF INTERESTS

The authors declare no competing interests.

---

**Figure S1.**
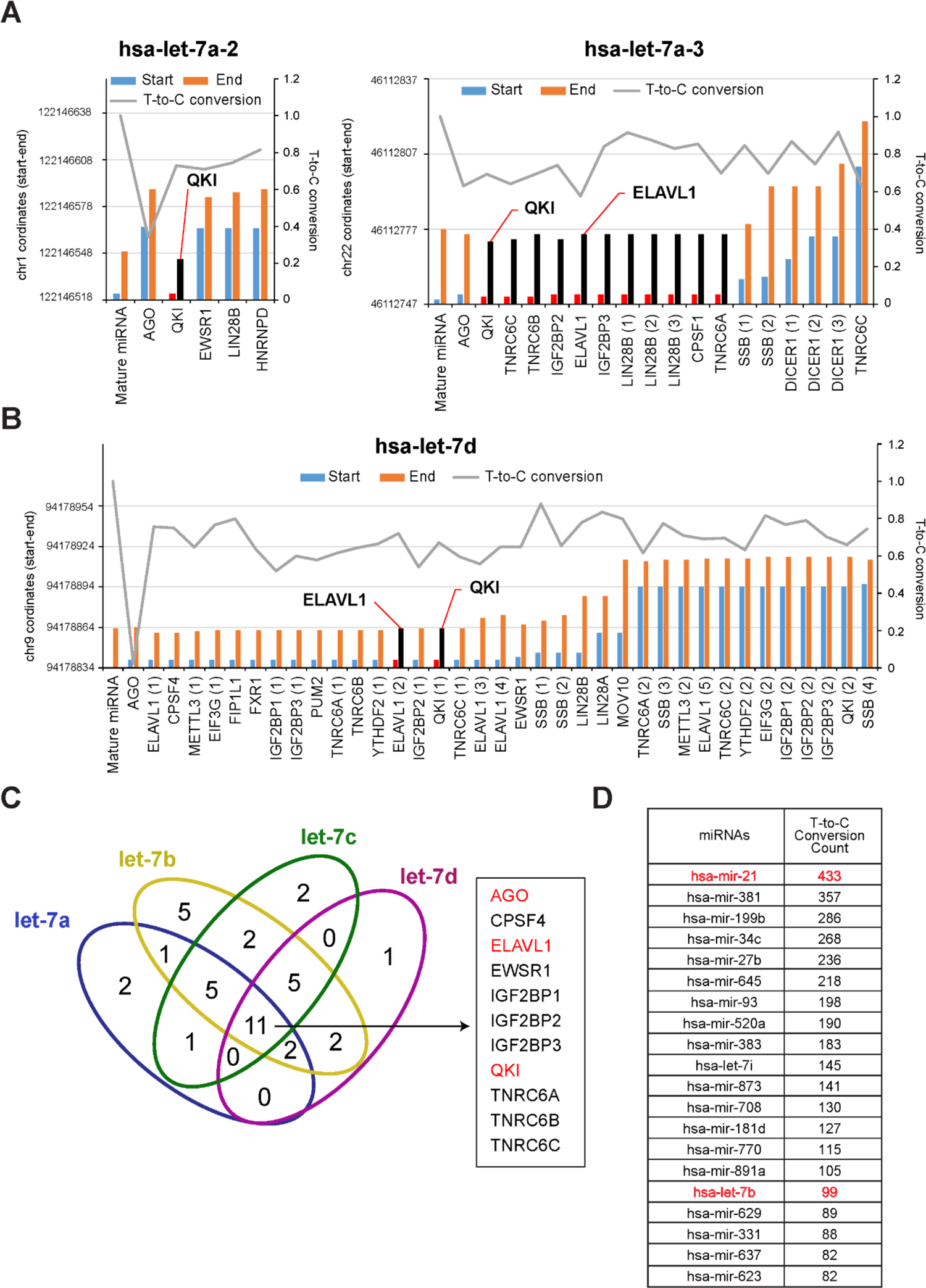
PAR-CLIP analysis identified RNA-biding proteins directly binding with mature and precursor let-7. **(A, B)** Genome coordinates (y-axis) of precursor and mature miRNA let-7a-2 (A, *Left*), 7a-3 (A, *Right*), and 7d (B) identified from PAR-CLIP using human cell lines. T-to-C conversion ratio was presented in y-axis as well. Representative RBPs such as QKI and HuR were highlighted. **(C)** Venn diagram of common RBPs directly binding with let-7a, 7b, 7c, and 7d. **(D)** We analyzed human QKI PAR-CLIP by sorting miRNAs in the order of T-to-C conversion counts: the top 20 miRNAs are listed.

**Figure S2.**
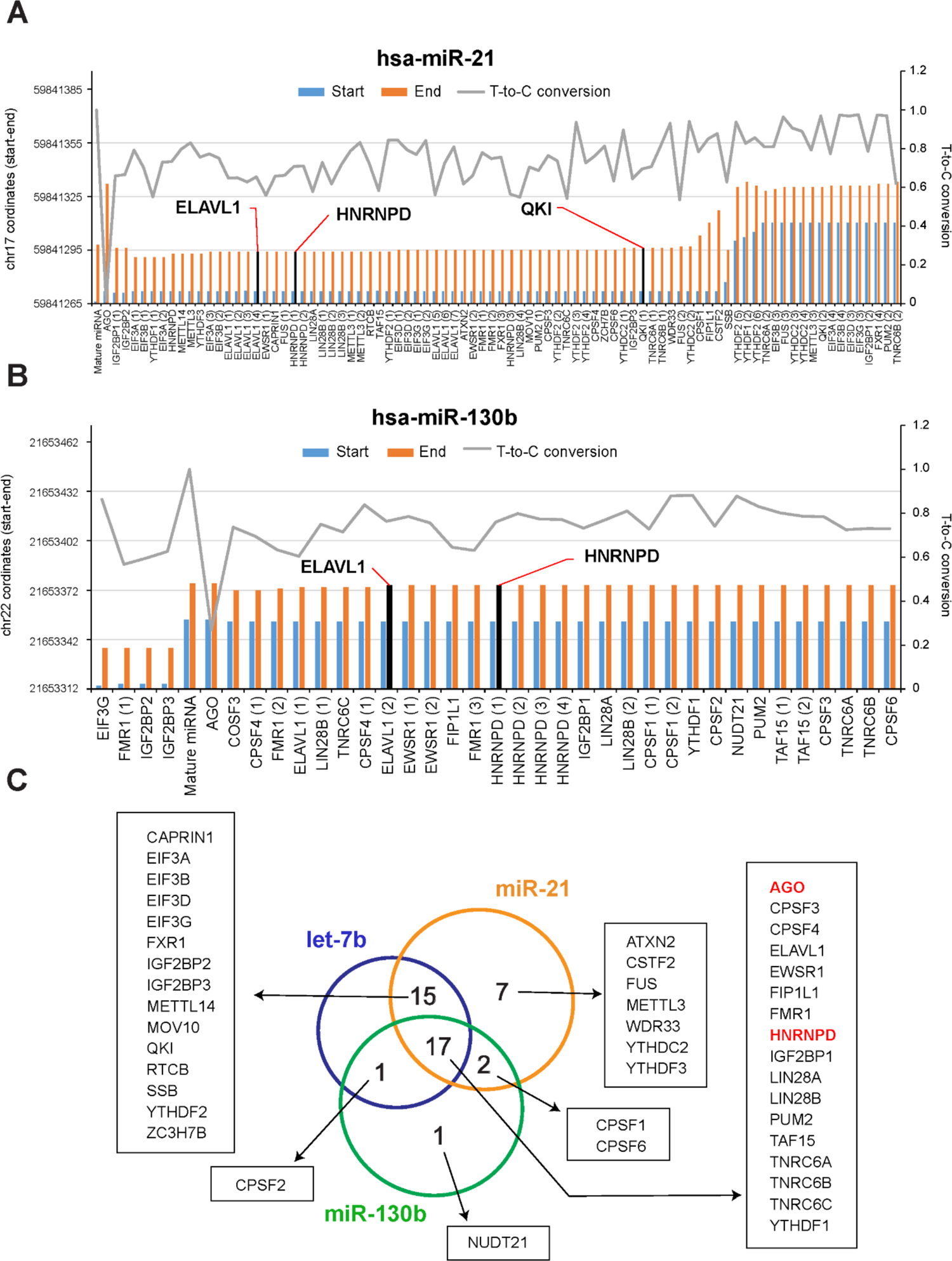
PAR-CLIP analysis identified RNA-biding proteins directly binding with mature and precursor miR-21 and miR-130b. **(A, B)** Genome coordinates (y-axis) of precursor and mature miRNA miR-21 (*Top*) and miR-130b (*Bottom*) identified from PAR-CLIP using human cell lines. T-to-C conversion ratio was presented in y-axis as well. Representative RBPs such as QKI, HuR and hnRNPD/AUF1 were highlighted. **(C)** Venn diagram of common RBPs directly binding with let-7b, miR-21, and miR-130b.

**Figure S3.**
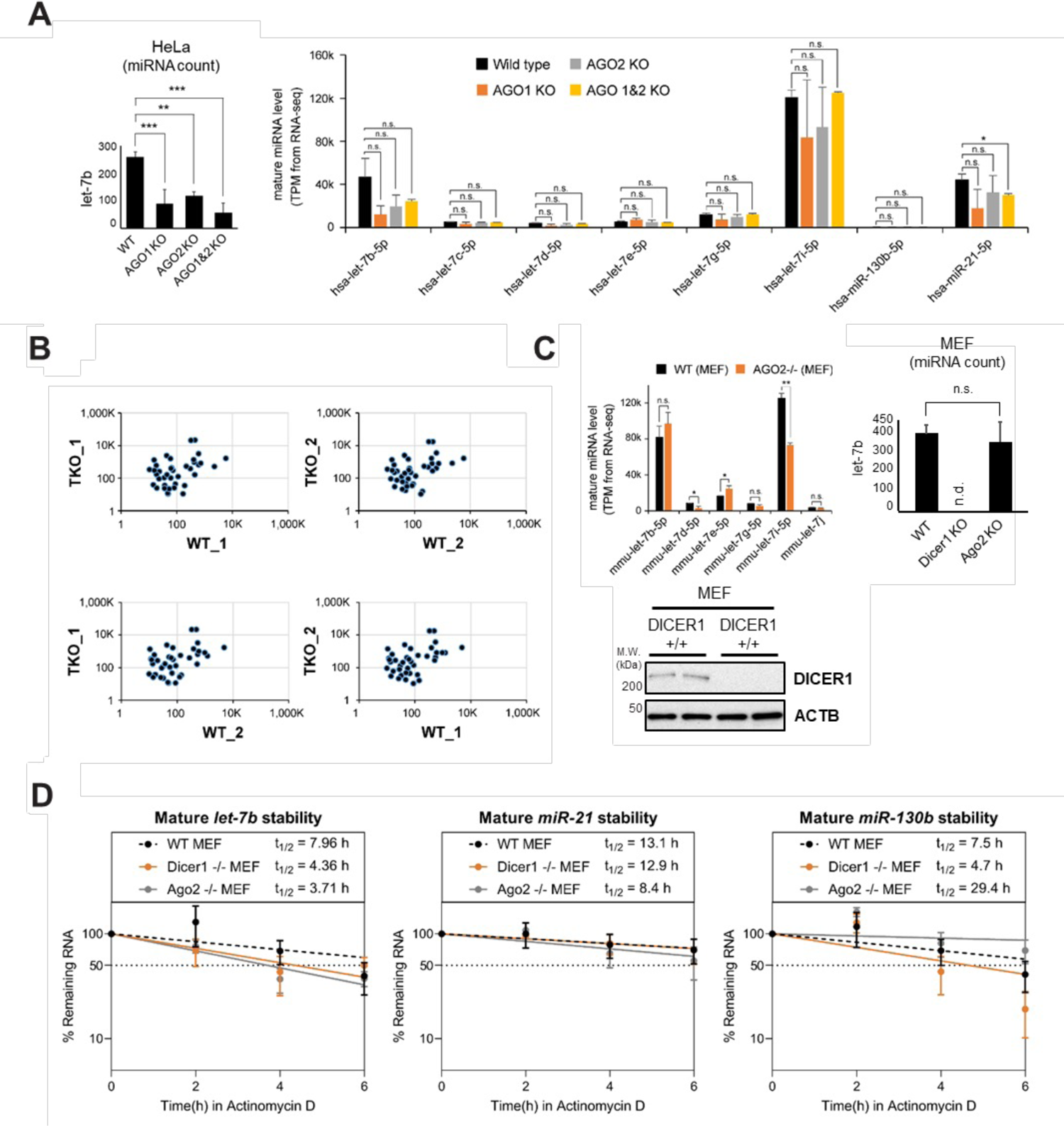
Assay of mature miRNAs when AGOs are depleted. **(A)** Level of let-7b was measured by miRNA counting (*Left*) and mature miRNA let-7, miR-130b and miR-21 levels were measured by RNA-seq (*Right*) using RNAs purified from wild type, AGO1 (Talen) KO, AGO2 (Talen) KO, or AGO1/2 DKO (Talen/CRISPR) HeLa cells. N = 2, \**p* < 0.05, *n.s.*, not significant from Student’s *t*-test. Data are mean ± S.D. **(B)** Scatter plots of passenger strand miRNAs from RNA-seq of HCT116 wild type and AGO TKO cells. **(C)** Levels of let-7 miRNAs were measured by RNA-seq (*Left*) and by miRNA counting (*Right*) using wild type, Dicer1 KO or Ago2 KO MEFs. (*Bottom*) Western blot analysis of Dicer1 and ACTB using MEF lysates. n.d., not detected from RNA-seq, \*\**p* < 0.01, \**p* < 0.05, *n.s.*, not significant from Student’s *t*-test. Data are mean ± S.D. **(D)** Stability of let-7b (*Left*), miR-21 (*Middle*), and miR-130b (*Right*) calculated from RT-qPCR using RNAs purified from wild type, Ago2 KO or Dicer1 KO MEFs treated with Actinomycin D at a given time point.

**Figure S4.**
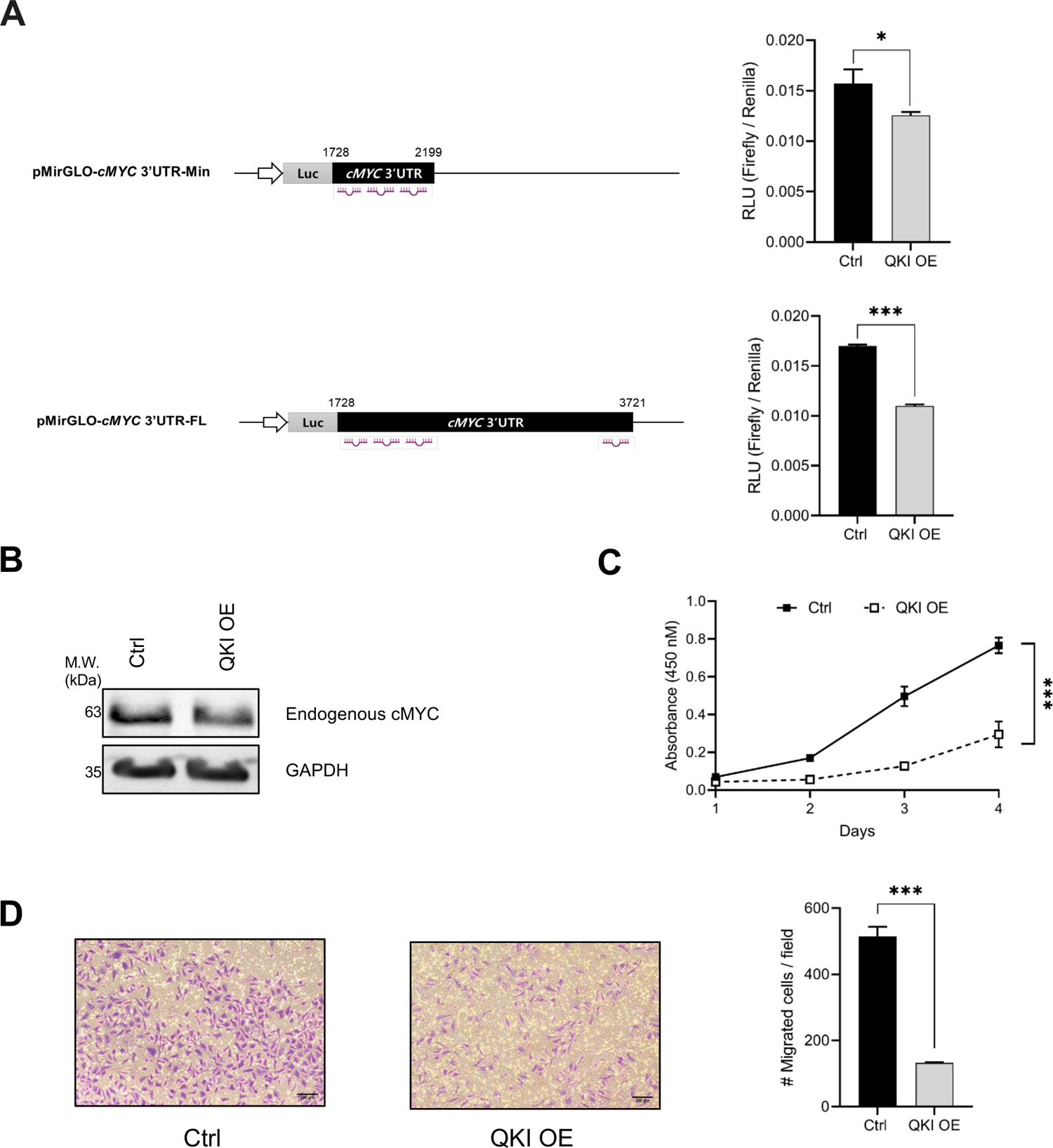
QKI suppresses proliferation and migration of cancer cell by enhancing 1et-7b activity. **(A)** Luciferase reporter analysis for let-7b activity after transfected with QKI overexpression plasmid in HeLa cells. N = 3, \*\*\**p < 0.001*, \**p < 0.05* from Student’s *t*-test. Data are mean ± S.D. **(B)** Western blot analyses of endogenous cMYC protein expression in HeLa cells transfected with either control or QKI overexpression plasmids. **(C)** Proliferation assay of HeLa cells that transfected with QKI overexpression plasmid. N = 4, *\*\*\*p < 0.001* from Student’s *t*-test. Data are mean ± S.D. **(D)** Migration assay in HeLa cells after QKI overexpression. N = 3, *\*\*\*p < 0.001* from Student’s *t*-test. Data are mean ± S.D.

**Figure S5.**
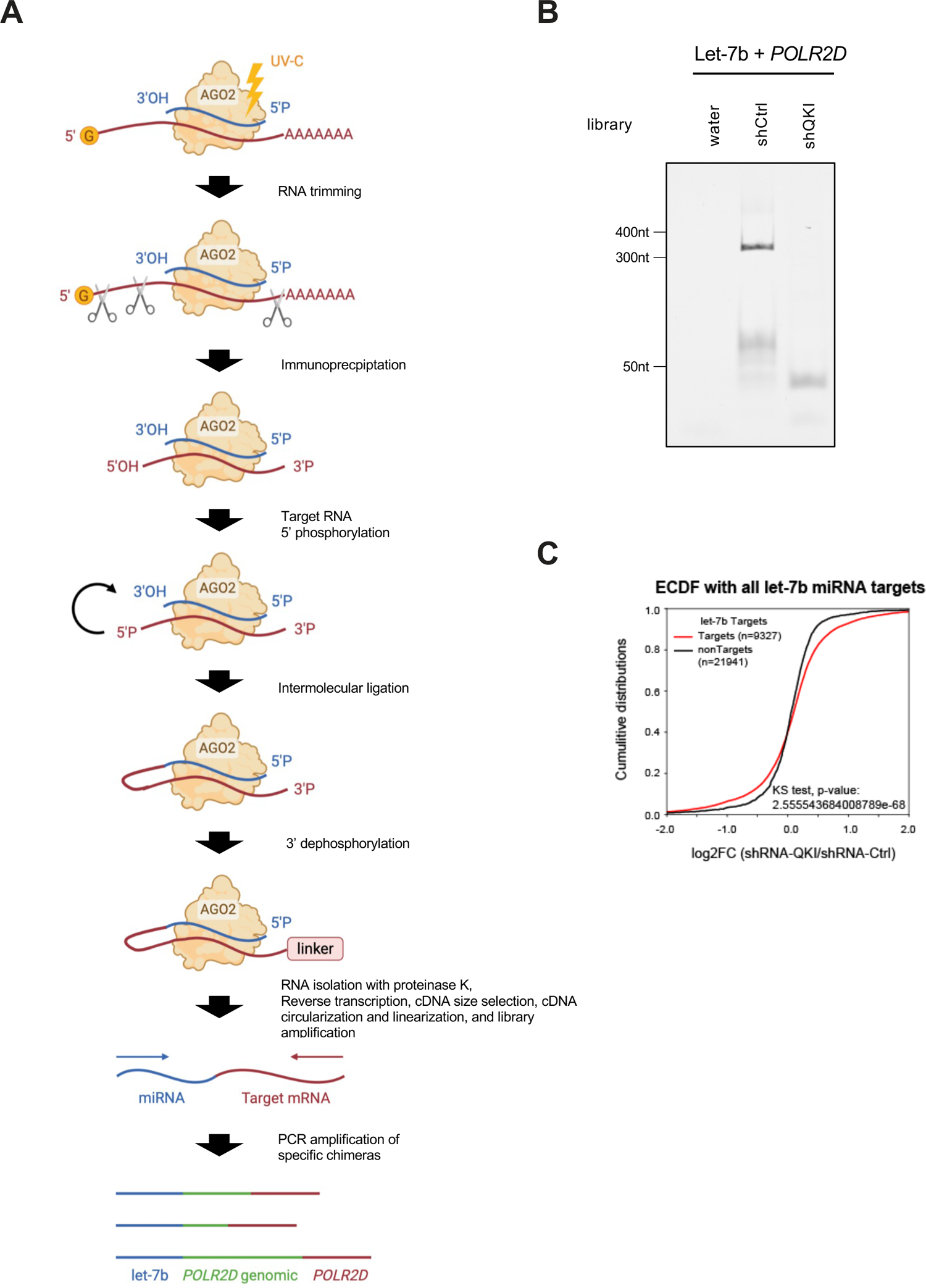
QKI stabilizes let-7b-*POLR2D* mRNA interaction. **(A)** Schematic representation of the main steps of Chimera PCR. **(B)** Detection of let-7b binding to the 3’ UTR of *POLR2D* mRNA using Chimera PCR after transfection of either control or QKI shRNAs in HeLa cells. **(C)** Cumulative-distribution fraction plots (CDF) depicting the let-7b target RNAs fold change upon QKI knockdown. Significance was determined using a two-sided Kolmogorov–Smirnov (KS) test.

